# Association of stress-related neural activity and baseline interleukin-6 plasma levels in healthy adults

**DOI:** 10.1101/2021.03.26.437134

**Authors:** Johanna F. Voges, Laura Müller-Pinzler, Miriam Neis, Finn Luebber, Tanja Lange, Jennifer E. Hundt, Meike Kasten, Ulrike M. Krämer, Sören Krach, Lena Rademacher

## Abstract

Several studies suggest a link between acute changes in inflammatory parameters due to an endotoxin or (psychological) stressor and the brain’s stress response. The extent to which basal circulating levels of inflammatory markers are associated with the brain’s stress response has been hardly investigated so far. In the present study, baseline plasma levels of the cytokine interleukin (IL)-6 were obtained and linked to neural markers of psychosocial stress using a modified version of the Montreal Imaging Stress Task in a sample of N=65 healthy subjects (N=39 female). Of three a-priori defined regions of interest – the amygdala, anterior insula, and anterior cingulate cortex – baseline IL-6 was significantly and negatively associated with stress-related neural activation in the right amygdala and left anterior insula. Our results suggest that baseline cytokines might be related to differences in the neural stress response and that this relationship could be inverse to that previously reported for induced acute changes in inflammation markers.

## 1. Introduction

Research has shown that peripheral cytokines can act on the brain via humoral, neural, and cellular pathways (see for example Miller et al. (2013) for review). This way, they interact with a variety of neural processes such as synaptic plasticity or neurotransmitter function and thereby affect behavior (Miller et al., 2009; Salvador et al., 2021). A specific profile of cytokine associations with neural processes in humans is, however, still unknown.

Eisenberger and colleagues suggested that pro-inflammatory cytokines affect social behavior, for example by increasing the brain’s threat-related sensitivity to negative social experiences (Eisenberger et al., 2017). A few fMRI studies approached this issue with paradigms in which unpleasant situations (negative social feedback, presentation of fearful faces, or a game simulating social exclusion) were induced. These studies either compared neural activity between a manipulation with bacterial endotoxins and placebo or correlated it to acute increases of inflammatory parameters (mainly interleukin (IL)-6) by endotoxin, stress, or the paradigm itself (Eisenberger et al., 2009; Inagaki et al., 2015; Muscatell et al., 2016, 2015; Slavich et al., 2010). They reported a positive relationship of acute inflammation with neural responses to negative social experiences, particularly in the amygdala, anterior cingulate cortex (ACC), dorsomedial prefrontal cortex (dmPFC), and anterior insula (AI). However, the association with basal circulating levels of inflammatory markers has been hardly investigated so far. Two previous studies linked baseline levels of pro-inflammatory (IL-6, IL-1 beta) and anti-inflammatory cytokines (IL-2) to neural activity during a social exclusion paradigm and found no significant association or even a negative correlation with pro-inflammatory and positive association with anti-inflammatory cytokines (Conejero et al., 2019; Slavich et al., 2010). Conejero et al. (2019) thus hypothesized that basal levels of peripheral cytokines and neural activation in response to social threats might compensate one another maintaining a state of homeostasis.

The current study addresses this issue by examining baseline blood levels of IL-6 and neural responses during acute social stress. Following the theory of Conejero and colleagues, we expected to find a negative relationship between IL-6 levels and neural stress responses. However, because those studies that had examined acute inflammatory responses reported positive correlations with neural activity (in line with the theory of Eisenberger and colleagues), we also wanted to explore whether the association was positive. The amygdala, AI, and ACC were defined as regions of interest as they were found to be related to both acute changes in inflammatory parameters in previous studies on neural responses to negative social experiences and to neurobehavioral responses to acute stress (see van Oort et al. (2017) for review). Furthermore, they are part of the interoceptive pathways which communicate peripheral inflammation to the brain (Savitz and Harrison, 2018). To further explore the association between basal IL-6 levels and stress reactions as well as chronic stress, we also assessed the acute cortisol and affective reaction to stress induction and obtained questionnaire measures of chronic stress and dispositional stress reactivity.

## 2. Methods

### 2.1 Participants

Prior to the experiments, all participants gave informed and written consent. The study was approved by the local ethics committee at the University of Lübeck (AZ 13-159). Participants were recruited through local advertisement and contact addresses provided by the residents’ registration office. As the current study was part of an interdisciplinary project which also examined ingestive behavior and obesity, participants had to meet the following inclusion criteria: scanner compatibility, no metabolic disorders, no current episode of major depression, mania or schizophrenia, no eating disorder, no alcohol misuse classified as ≥ 8 points in the alcohol use disorder identification test (Saunders et al., 1993), and no intake of antidepressants, antipsychotics, antihistamines, beta blockers, corticosteroids, valproate, lithium, antiemetics, laxatives, amphetamines, or other stimulants.

From the sample of the interdisciplinary project (N=184, 108 female), only participants with complete fMRI scans who provided plasma samples were included in the present analyses (N=86). Unfortunately, one plate of the IL-6 measurement did not meet the quality standards in terms of variation to the other two plates (inter-assay coefficient of variation > 15%), so the values of this plate were not included in the analyses (N=21). The resulting sample consisted of N= 65 participants (39 female). Participants’ age ranged from 28 to 54 years (mean age 42.6 ± 7.47 years), the mean Body Mass Index (BMI) was 26.4 ± 4.95 (more demographic information can be found in S1). For questionnaire measures, there were missing values (see S2 for details) so that analyses on these parameters were performed for slightly smaller sample sizes (N= 63 - 64).

### 2.2 Procedure

The study took place on two different days and consisted of several experiments that are not all part of this analysis (e.g., tests on eating behavior). On one of the days, fasting blood samples were drawn and body fat was determined. This was always scheduled in the morning (between 8 am and 10.30 am) to reduce circadian influences. On the other day (the order differed between participants), psychosocial stress was induced during fMRI scanning. For organizational reasons, this investigation could not always be performed at the same time of day. Before and immediately after stress induction, saliva samples were obtained for cortisol measurement and participants filled in the Positive and Negative Affect Schedule (PANAS; Watson et al., 1988) as a measure of the affective response to stress. Furthermore, participants completed the Trier Inventory for Chronic Stress (TICS; Schulz et al., 2004) and the Stress Reactivity Scale (SRS; Schulz et al., 2005) to obtain measures for chronic stress in the last three months and stress reactivity in the sense of a disposition that underlies individual differences in stress reaction (see below for details).

### 2.3 Stress induction

For stress induction, a modified version of the Montreal Imaging Stress Task (MIST) by Dedovic and colleagues was used (Dedovic et al., 2005). Participants were supposed to solve mental arithmetical problems of varying difficulty. The task was performed in three runs, in which two blocks of stress conditions alternated with a control condition. In the stress conditions, the time limit for solving the questions was adapted to the participant’s performance in order to reduce the success rate in comparison to the control condition. A bar on the screen visualized the time that had already elapsed while processing the tasks. Arrows indicated the participant’s own performance in comparison to a reference group (please see supplementary section S3 for an example of the screen). To increase pressure, the experimenter provided negative feedback on the participants’ performance by telling them via speaker: ‘You unfortunately still make many mistakes.’ If a participant solved a row of tasks correctly, the feedback was reformulated: ‘Unfortunately you are not fast enough.’ Time constraints of the task and especially the social evaluative component were supposed to induce psychosocial stress in the stress condition (Dickerson and Kemeny, 2004). During the control condition, there was no time limit and no feedback on task performance. The two blocks of stress condition were initially planned to consider differences between two distinct stress conditions: In one condition, a frame around the screen indicated that an audience of three persons of the study staff was watching the participant’s performance live. In the other condition, this component was missing. Since we did not assume that this experimental manipulation would affect the research question followed in this publication, data of these two conditions were collapsed. The experiment was run in a fixed order and lasted on average 25.7 minutes. Before scanning, the participants practiced the task in order to become familiar with the procedure.

Two parameters were used as behavioral measures of performance: The mean reaction time required to correctly solve the arithmetic tasks and the percentage of correctly solved tasks. To investigate stress-related changes in performance, we compared the mean of the control condition to the mean of the stress condition for both reaction time and percentage of correctly answered tasks. There was a significant decrease from control to stress condition in reaction time (t(64) = 16.7, p < 0.001, d= 2.08) and correctly answered tasks (t(64) = 20.5, p< 0.001, d= 2.54, see also Table 1).

**Table 1.**
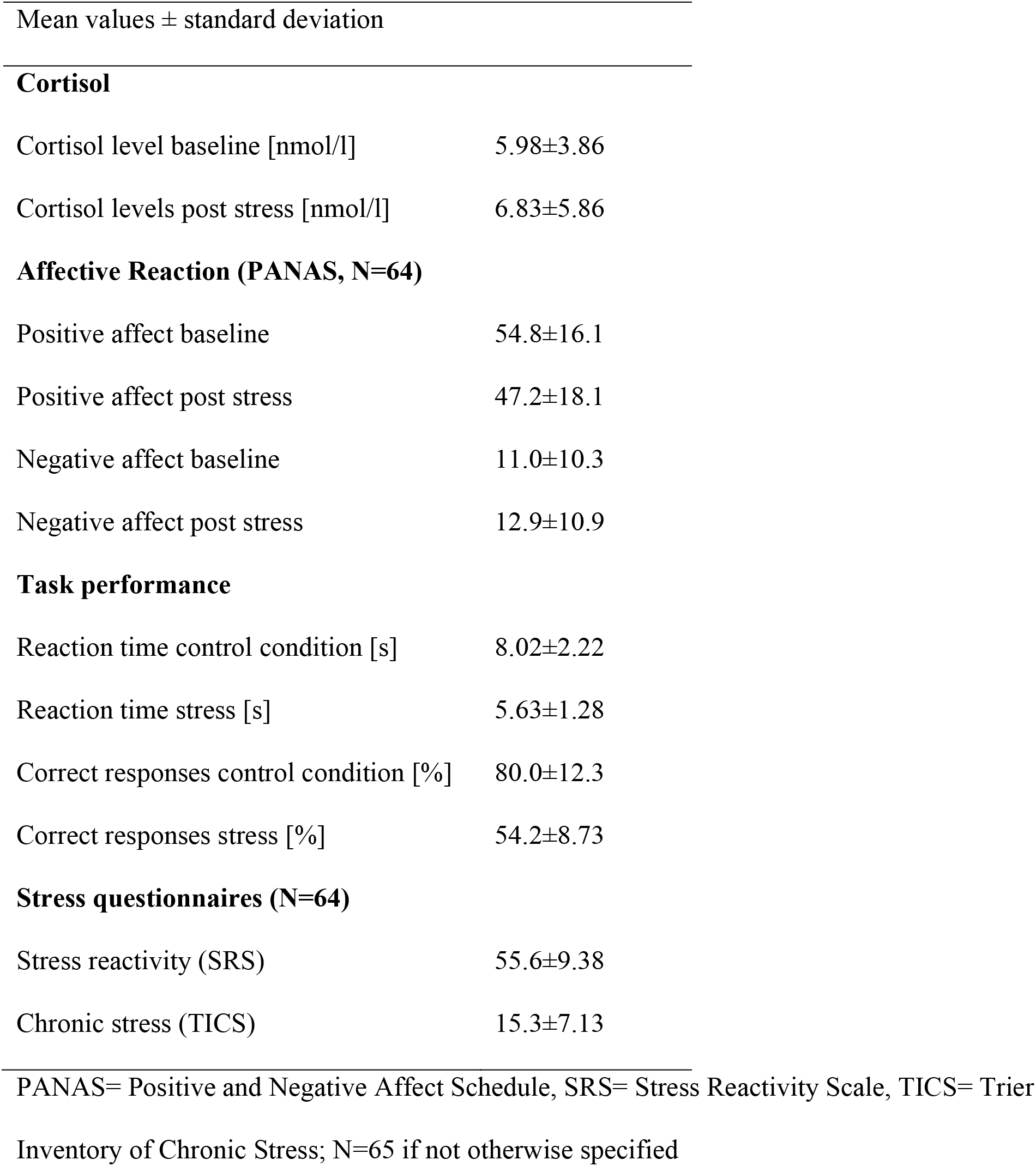
Descriptive statistics of acute stress task response and stress questionnaires

| Mean values ± standard deviation | |
| --- | --- |
| **Cortisol** |  |
| Cortisol level baseline [nmol/l] | 5.98±3.86 |
| Cortisol levels post stress [nmol/l] | 6.83±5.86 |
| **Affective Reaction (PANAS, N=64)** |  |
| Positive affect baseline | 54.8±16.1 |
| Positive affect post stress | 47.2±18.1 |
| Negative affect baseline | 11.0±10.3 |
| Negative affect post stress | 12.9±10.9 |
| **Task performance** |  |
| Reaction time control condition [s] | 8.02±2.22 |
| Reaction time stress [s] | 5.63±1.28 |
| Correct responses control condition [%] | 80.0±12.3 |
| Correct responses stress [%] | 54.2±8.73 |
| **Stress questionnaires (N=64)** |  |
| Stress reactivity (SRS) | 55.6±9.38 |
| Chronic stress (TICS) | 15.3±7.13 |
| PANAS= Positive and Negative Affect Schedule, SRS= Stress Reactivity Scale, TICS= Trier Inventory of Chronic Stress; N=65 if not otherwise specified |  |

### 2.4 Questionnaires

To assess changes in mood during the experiment, measures of positive and negative affect were obtained before and after stress induction using the PANAS (Watson et al., 1988). The PANAS comprises ten positive and ten negative adjectives to describe the current affective state on a five-point scale. Positive and negative affect obtained before stress induction was subtracted from the score after scanning to get a measure of affective response to the MIST. To assess perceived chronic stress during the last three months, participants filled out the TICS (Schulz et al., 2004). The questionnaire consists of 57 items with subscales for different types of stress and is answered on a five-point Likert scale. The 12-item screening scale of the TICS was used as a global measure for experienced stress.

Dispositional stress reactivity was measured with the SRS that consists of 29 items with three response options (Schulz et al., 2005). Stress reactivity is herein defined as a constitutional trait that relatively stable constitutes the individual reaction to stressors (Schlotz et al., 2011).

### 2.5 Cortisol

Cortisol was assessed from saliva samples collected one hour before the stress induction (30 minutes after the participant’s arrival) and approximately 5 minutes after scanning (average time after onset of the stress task = 30.6 ± 2.9 minutes). Saliva samples were frozen and stored at −20 °C until they were sent to the lab of Dresden LabService GmbH. After thawing, salivettes were centrifuged at 3000 rpm for five minutes, which resulted in a clear supernatant of low viscosity. Salivary concentrations were measured by commercially available chemiluminescence immunoassay with high sensitivity (IBL International, Hamburg, Germany).

### 2.6 IL-6

Blood samples were obtained by antecubital venipuncture, collected in EDTA tubes (5 ml) and placed on ice. Immediately after collection, the samples were centrifuged at 2000 g for ten minutes at 4 °C and plasma was then stored at −80 °C until further analysis. Interleukin-6 was measured in duplicate using a commercially available high-sensitive enzyme-linked immunosorbent assay (ELISA, R&D Systems, Minneapolis) with a minimum detection threshold of 0.2 pg/ml. Within-assay coefficients of variation were <9%, the between-assay coefficient was 13.6%. IL-6 data were positively skewed, so raw values were log transformed to normalize the distribution prior to statistical testing.

### 2.7 fMRI scanning and data processing

Participants were scanned on a 3 Tesla MAGNETOM Skyra MR tomograph (Siemens) equipped with a 64-channel head-coil at the Center of Brain, Behavior and Metabolism in Lübeck, Germany. A BOLD sensitive echo planar imaging (EPI) sequence was used for acquisition of functional volumes during the experiment (TR = 1.5 s, TE = 25.0 ms, flip angle = 80°, 56 interleaved/ascending slices, slice thickness = 3 mm, FOV= 512×512). The number of scans varied between the participants as they needed different amounts of time for the task (mean per run= 309 scans ±22). Additionally, structural images of the whole brain were acquired (TR = 1.9s, TE = 2.44, flip angle = 9°, slice thickness = 1 mm, FOV = 256×256). Functional MRI data analysis was performed using SPM12 (Wellcome Department of Imaging Neuroscience, London, UK; http://www.fil.ion.ucl.ac.uk/spm) implemented in Matlab R2019b (Mathworks Inc., Sherborn, MA, USA). Functional images were slice timed and realigned to the mean image to correct for head motions. Next, they were normalized into MNI space using the standard segmentation as implemented in SPM12 on the mean functional image and tissue probability maps. Normalization parameters were then applied on all images of the time series. Finally, functional images were spatially smoothed with an 8 mm full width half maximum isotropic Gaussian kernel and high-pass filtered at 1/256 Hz to remove low frequency drifts.

### 2.8 Statistical analysis of fMRI data

Statistical analysis of the fMRI data was performed in a two-level, mixed-effects procedure. The generalized linear model (GLM) on the first level included regressors for the three conditions (control condition, two stress conditions) and the six motion parameters as regressors of no interest. Linear contrasts were computed for each participant that compared blood oxygenation level dependent (BOLD) signal during stress to the control condition. These contrast images of each participant were then used for random effects analyses at the group level. First, two one-sample t-tests were calculated to assess stress task-related neural activation (stress conditions > control condition) and deactivation (control condition > stress condition). Second, to investigate whether IL-6 levels were associated with stress-related neural activation, IL-6 was included as a covariate in the one-sample t-test of stress task-related activation and positive as well as negative correlation contrasts were calculated.

Region-of-interest (ROI) analyses were performed in the amygdala, insular cortex, and ACC for the contrasts mentioned above. ROIs for the bilateral amygdala and the ACC were defined anatomically based on the Automated Anatomical Labeling Atlas (Tzourio-Mazoyer et al., 2002) with a dilation factor of one. The bilateral anterior insula ROI was based on the two anterior insula clusters from the three cluster solution by DiMartino and Kelly (Kelly et al., 2012).

If not mentioned otherwise, all results were corrected for multiple comparisons applying family-wise error (FWE) correction as implemented in SPM12. All results are reported in MNI space.

### 2.9 Statistical analyses of stress parameters

Statistical analyses were performed using the open source software Jamovi (The jamovi project, 2021). First, to investigate whether the MIST task induced stress in the current sample, one-tailed Student’s t-tests were computed to test for changes in cortisol as well as negative and positive affect (before to after stress induction). Next, we investigated whether IL-6 was related to the acute cortisol and affective reaction to stress as well as chronic stress levels and stress reactivity. To this end, we tested for associations between IL-6 and change in cortisol, negative and positive affect as well as stress reactivity and chronic stress using Pearson correlations. The time of the day of cortisol collection was included as covariate in the correlation analysis involving cortisol, because cortisol plasma levels have a circadian rhythm (Lee et al., 2015). All reported results were considered to be significant at the p <0.05 level.

## 3. Results

### 3.1. Stress task effects

On average, cortisol levels increased from before to after stress induction by 29.8% (see Table 1). However, this increase was not significant (t(64) = −1.13, p = 0.132, d= −0.14) due to great individual differences: only 43.1% of the participants showed increases in their cortisol levels (‘cortisol responders’), the remaining 56.9% showed no alterations or even a decline in cortisol levels. The observed percentage of cortisol responders is similar to previous studies using the MIST (Khalili-Mahani et al., 2010; Wheelock et al., 2016). Positive affect significantly decreased after stress induction (t(62) = 5.61, p< 0.001, d= 0.71), but changes in negative affect did not reach significance (t(62) = −1.60, p = 0.057, d= 0.20, see also Table 1). A decrease in positive affect was seen in 73.0 % of participants, an increase in negative affect in 49.2%.

### 3.2. Association of IL-6 with chronic stress levels and stress reactivity

IL-6 did not correlate with questionnaire measures of stress reactivity (r = −0.05, p= 0.70), chronic stress (r = −0.02, p= 0.88), or change of affect or cortisol after stress induction (positive affect: r =.06, p= 0.65; negative affect: r =.05, p= 0.71; cortisol: r = −0.17, p = 0.19).

### 3.3. Neural correlates of the stress task

When comparing stress to the control condition of the MIST (stress > control condition), increased brain activity was found in the bilateral middle temporal gyrus, superior medial and inferior orbital frontal gyrus, cuneus, cerebellum, temporal pole, right insular cortex, and mid cingulum (FWE corrected for multiple comparisons; see Table 2 and S4). In addition, analyses within our previously specified ROIs revealed significant effects in the bilateral amygdala and ACC and bilateral anterior insula (see Table 2). The opposite contrast (control condition > stress) revealed activation in the superior parietal and inferior occipital lobule, angular gyrus and supramarginal gyrus (see Table 3 and S4). In the previously specified ROIs no significant effects were found for this contrast.

**Table 2.**
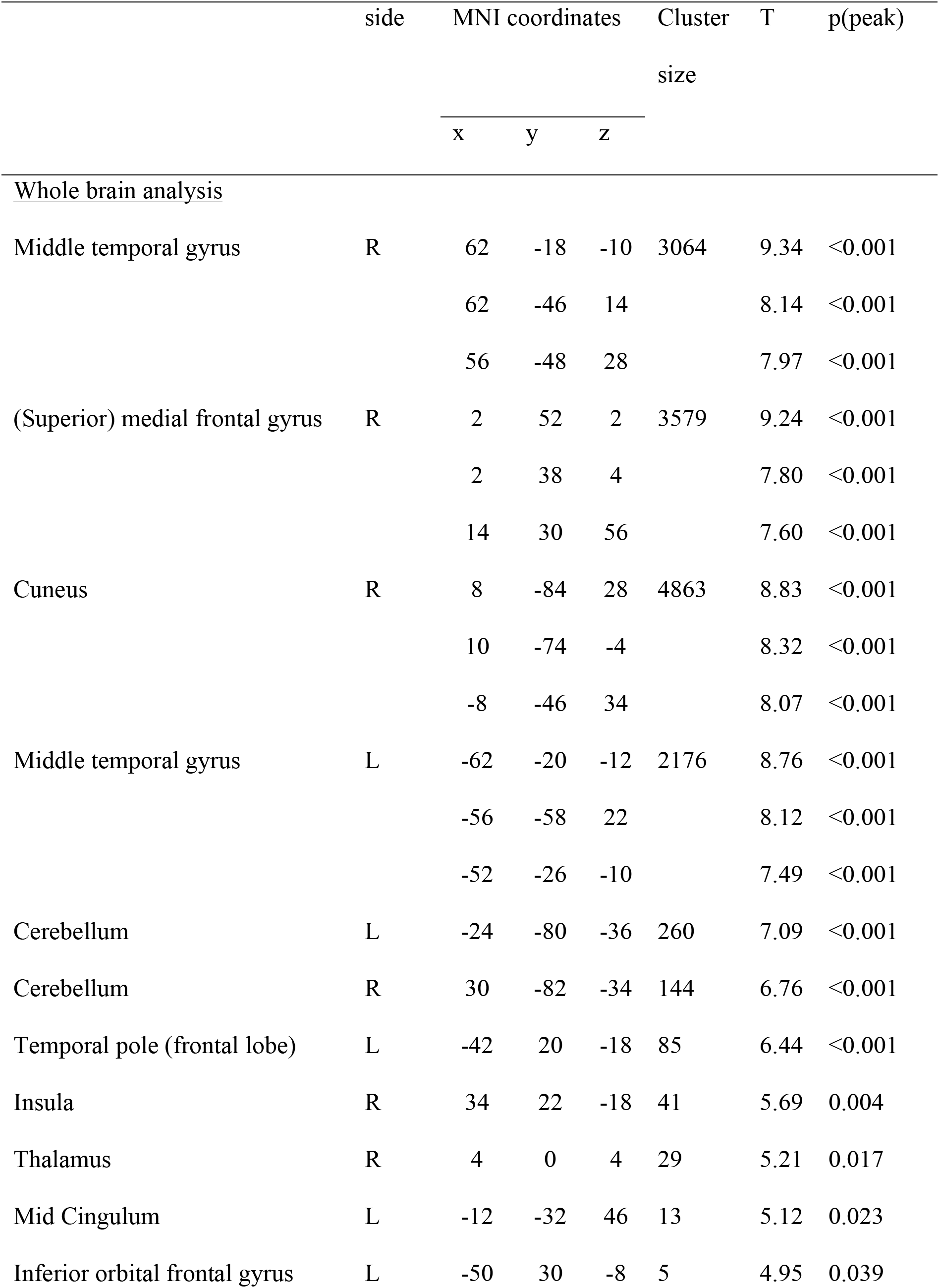
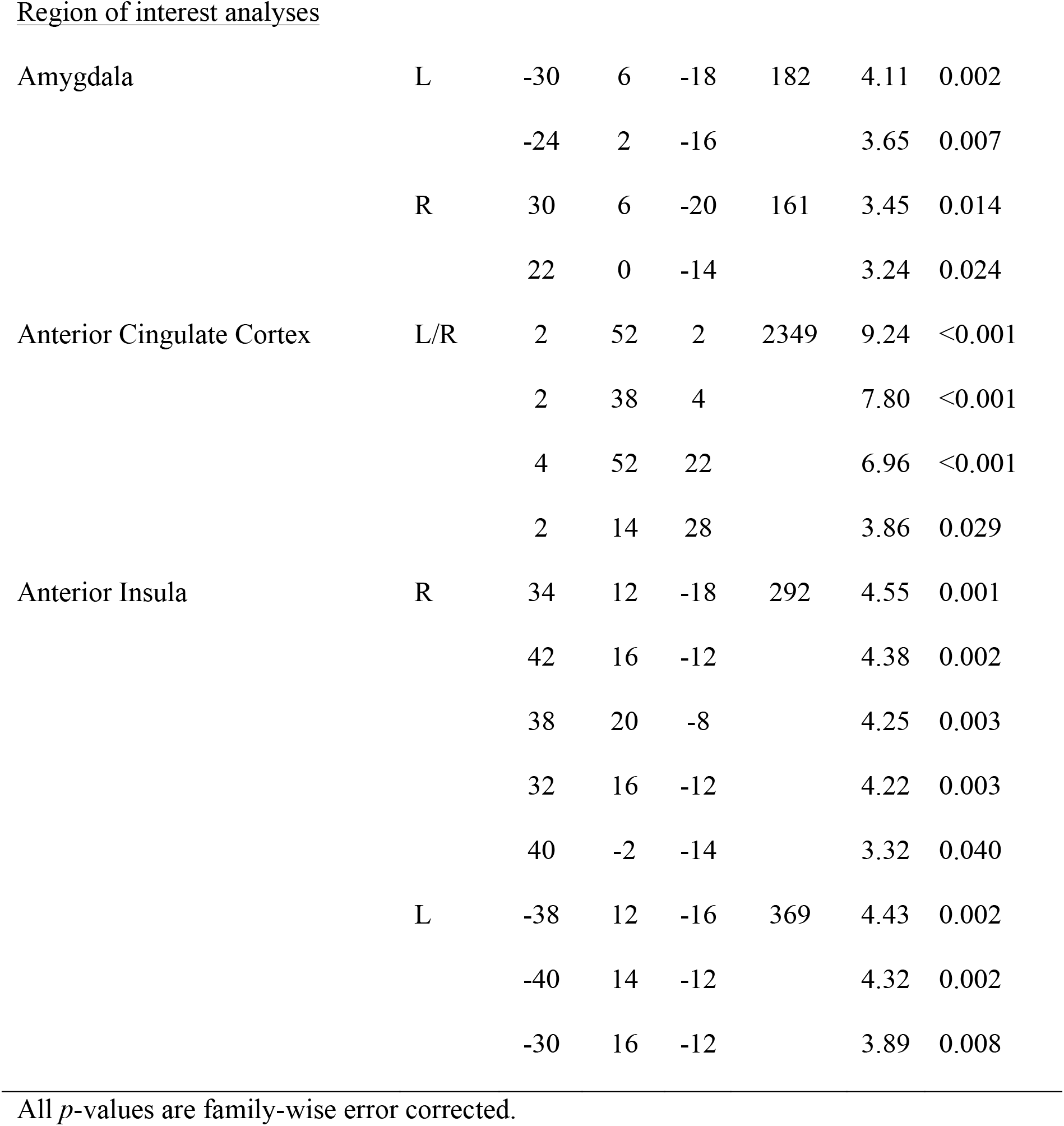
Activation at contrast stress > control condition

|  | side | MNI coordinates |  |  | Cluster<br>size | T | p(peak) |
| --- | --- | --- | --- | --- | --- | --- | --- |
|  |  | x | y | z |  |  |  |
| <u>Whole brain analysis</u> |  |  |  |  |  |  |  |
| Middle temporal gyrus | R | 62 | -18 | -10 | 3064 | 9.34 | <0.001 |
|  |  | 62 | -46 | 14 |  | 8.14 | <0.001 |
|  |  | 56 | -48 | 28 |  | 7.97 | <0.001 |
| (Superior) medial frontal gyrus | R | 2 | 52 | 2 | 3579 | 9.24 | <0.001 |
|  |  | 2 | 38 | 4 |  | 7.80 | <0.001 |
|  |  | 14 | 30 | 56 |  | 7.60 | <0.001 |
| Cuneus | R | 8 | -84 | 28 | 4863 | 8.83 | <0.001 |
|  |  | 10 | -74 | -4 |  | 8.32 | <0.001 |
|  |  | -8 | -46 | 34 |  | 8.07 | <0.001 |
| Middle temporal gyrus | L | -62 | -20 | -12 | 2176 | 8.76 | <0.001 |
|  |  | -56 | -58 | 22 |  | 8.12 | <0.001 |
|  |  | -52 | -26 | -10 |  | 7.49 | <0.001 |
| Cerebellum | L | -24 | -80 | -36 | 260 | 7.09 | <0.001 |
| Cerebellum | R | 30 | -82 | -34 | 144 | 6.76 | <0.001 |
| Temporal pole (frontal lobe) | L | -42 | 20 | -18 | 85 | 6.44 | <0.001 |
| Insula | R | 34 | 22 | -18 | 41 | 5.69 | 0.004 |
| Thalamus | R | 4 | 0 | 4 | 29 | 5.21 | 0.017 |
| Mid Cingulum | L | -12 | -32 | 46 | 13 | 5.12 | 0.023 |
| Inferior orbital frontal gyrus | L | -50 | 30 | -8 | 5 | 4.95 | 0.039 |
| Region of interest analyses |  |  |  |  |  |  |  |
| Amygdala | L | -30 | 6 | -18 | 182 | 4.11 | 0.002 |
|  |  | -24 | 2 | -16 |  | 3.65 | 0.007 |
|  | R | 30 | 6 | -20 | 161 | 3.45 | 0.014 |
|  |  | 22 | 0 | -14 |  | 3.24 | 0.024 |
| Anterior Cingulate Cortex | L/R | 2 | 52 | 2 | 2349 | 9.24 | <0.001 |
|  |  | 2 | 38 | 4 |  | 7.80 | <0.001 |
|  |  | 4 | 52 | 22 |  | 6.96 | <0.001 |
|  |  | 2 | 14 | 28 |  | 3.86 | 0.029 |
| Anterior Insula | R | 34 | 12 | -18 | 292 | 4.55 | 0.001 |
|  |  | 42 | 16 | -12 |  | 4.38 | 0.002 |
|  |  | 38 | 20 | -8 |  | 4.25 | 0.003 |
|  |  | 32 | 16 | -12 |  | 4.22 | 0.003 |
|  |  | 40 | -2 | -14 |  | 3.32 | 0.040 |
|  | L | -38 | 12 | -16 | 369 | 4.43 | 0.002 |
|  |  | -40 | 14 | -12 |  | 4.32 | 0.002 |
|  |  | -30 | 16 | -12 |  | 3.89 | 0.008 |

All *p*-values are family-wise error corrected.

**Table 3.** Activation at contrast control > stress (whole brain analysis)

|  | side | MNI coordinates |  |  | Cluster size | T | p(peak) |
| --- | --- | --- | --- | --- | --- | --- | --- |
|  |  | x | y | z |  |  |  |
| Superior parietal lobule | L | -26 | -60 | 52 | 1066 | 7.72 | <0.001 |
|  |  | -38 | -40 | 42 |  | 6.86 | <0.001 |
|  |  | -22 | -62 | 62 |  | 6.17 | <0.001 |
| Inferior occipital lobule | R | 34 | -88 | -2 | 244 | 7.41 | <0.001 |
|  |  | 28 | -94 | 4 |  | 6.48 | <0.001 |
| Angular gyrus | R | 28 | -60 | 50 | 417 | 6.94 | <0.001 |
|  |  | 30 | -52 | 50 |  | 6.26 | <0.001 |
| Inferior occipital lobule | L | -30 | -90 | -10 | 129 | 6.27 | <0.001 |
|  |  | -30 | -92 | 0 |  | 6.19 | <0.001 |
|  |  | -22 | -96 | 4 |  | 5.85 | <0.001 |
| Supramarginal gyrus | R | 42 | -36 | 46 | 61 | 5.87 | 0.002 |

All *p*-values are family-wise error corrected.

**Table 4.** Correlations of IL-6 with neural activation at contrast stress > control condition in regions of interest.

|  | side | MNI coordinates |  |  | Cluster | T | p (peak) |
| --- | --- | --- | --- | --- | --- | --- | --- |
|  |  |  |  |  | size |  |  |
|  |  | x | y | z |  |  |  |
| Amygdala | R | 34 | -4 | -28 | 5 | 3.14 | 0.030 |
|  |  | 28 | -6 | -16 | 6 | 2.97 | 0.046 |
| Anterior Insula | L | -40 | 4 | -16 | 3 | 3.41 | 0.029 |

All *p*-values are family-wise error corrected.

### 3.4. Association of stress task-related neural activity and IL-6

There was no positive association of stress task-associated neural activity (stress > control condition) with IL-6 levels, but ROI analyses revealed negative correlations with the activity of the left anterior insula and right amygdala (FWE corrected, see Figure 1 and Table 4, for results of whole brain analysis see S5).

**Figure 1.**
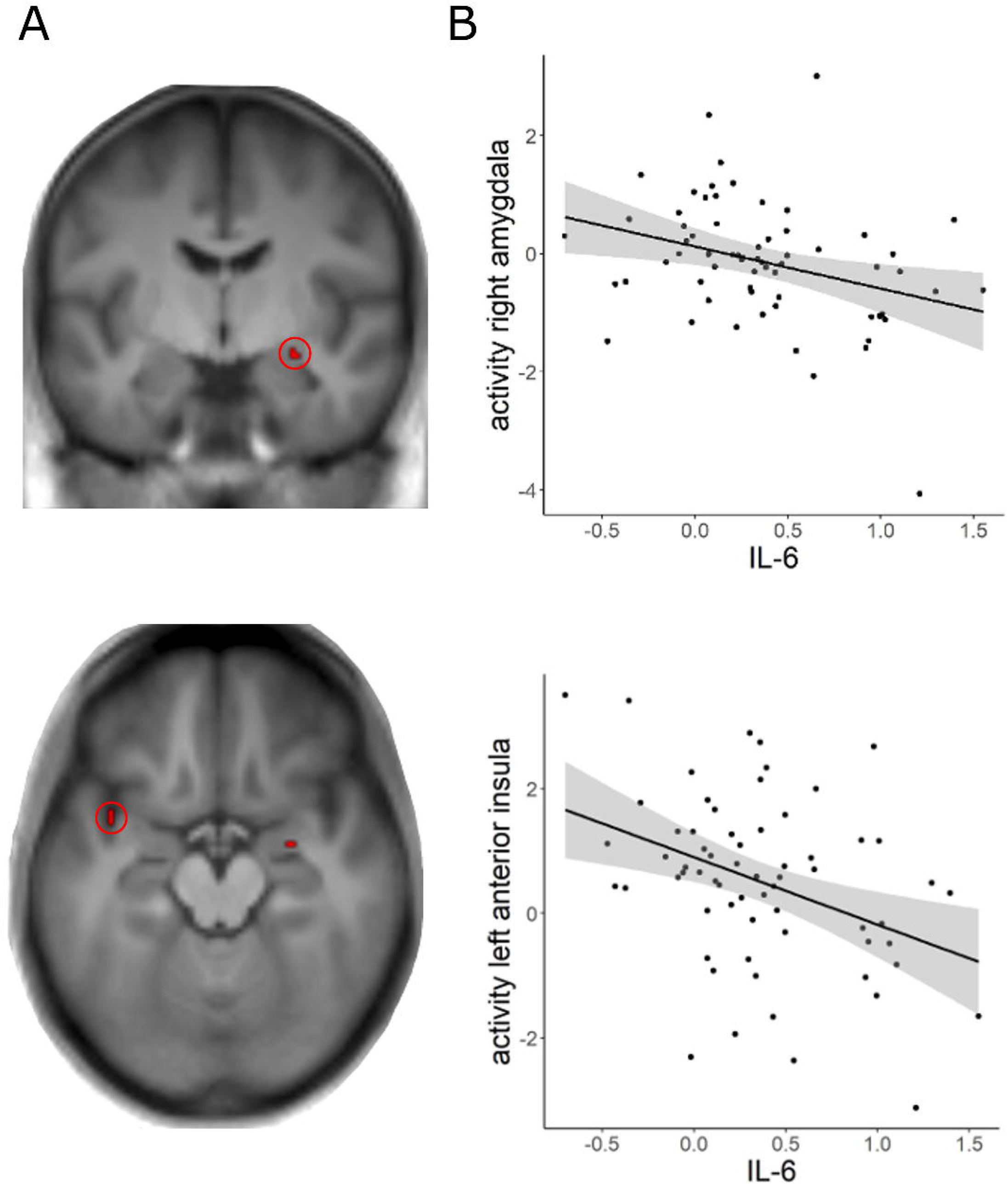
Correlation of log transformed baseline interleukin-6 (IL-6) levels and stress task-related neural activity A) Cluster of significant correlation between IL-6 and neural activity for the contrast stress > control condition in the right amygdala and left anterior insula (p=0.05, FWE corrected, clusters are displayed on the average T1 image of the sample). B) Scatter plot of the correlation between IL-6 and parameter estimates of the activation in the left anterior insula and right amygdala. For visualization purposes, parameter estimates were extracted from a sphere of 6 mm around the peak activation (anterior insula: −40, 4, −16; amygdala: 34, −4, −28). Alt Text for illustration [27 words]: On the left are two MRI images of the brain showing clusters of functional activity. On the right are two scatterplots with regression lines and confidence intervals.

## 4. Discussion

The aim of the present study was to investigate the association of baseline IL-6 levels and neural responses to psychosocial stress. For this purpose, blood was collected from healthy participants and neural activity during an fMRI stress paradigm was examined in three regions of interest - the amygdala, AI, and ACC. Negative associations between baseline IL-6 levels and activity in the left AI and right amygdala during stress exposure were found. These findings suggest that the relationship between basal IL-6 and social stress-related brain activity could be inverse to that previously reported for induced acute changes in IL-6 (Eisenberger et al., 2009; Muscatell et al., 2016, 2015). However, several limitations of the study, that render the interpretation of the findings difficult, are discussed below.

Regarding the activation of the amygdala and the anterior insula during stress exposure, our study replicates previous findings that compared evaluative stress to a non-stressful control condition (e.g., Chung et al., 2016; Lederbogen et al., 2011; Orem et al., 2019; Zschucke et al., 2015). However, there have also been studies reporting no stress-related effects in these regions or even deactivations and the reasons for this heterogeneity of results are still unclear (see van Oort et al. (2017) for review). Importantly, activation of the amygdala and AI have also been linked to the central processing of interoceptive signals (Chen et al., 2021). Interoception describes the processes by which the nervous system senses and integrates -and in a recently proposed definition also regulates - signals from the internal organs (Chen et al., 2021). The AI is assumed to integrate viscerosensory signals with representations of motivational, social and cognitive conditions (Craig, 2009). The amygdala is densely connected to the AI and is involved in affective processes (Adolphs et al., 1995; Morris et al., 1998), especially in an evaluative context (Guyer et al., 2008; Lorberbaum et al., 2004; Müller-Pinzler et al., 2015). The neurally mediated interoceptive pathways including the AI and amygdala have been shown to also become activated by peripheral inflammation (Savitz and Harrison, 2018), which could be related to the finding of an association between IL-6 levels and activity in the left AI and right amygdala in the current study.

The main finding of our study is that the association between baseline IL-6 levels and neural markers of stress response was negative. This result is in line with the theory of Conejero and colleagues (2019) that basal inflammation might prevent excessive neural activation in response to social threats. In their study the authors suggested that peripheral cytokines and brain activity could be regarded as parts of a social warning system interacting in a balanced relationship. However, Conejero and colleagues based their reasoning on the finding of a negative correlation of neural activity in the ACC during a social exclusion paradigm with pro-inflammatory cytokine IL-1 beta and a positive correlation of activation in the ACC, insula, and OFC with anti-inflammatory IL-2. For IL-6, neither Conejero et al. (2019) nor another study by Slavich and colleagues et al. (2010) found a significant association with neural activity. This disconnect between the studies so far needs more careful consideration: Neuroimaging research on the association of peripheral immune parameters and neural function has mostly reduced cytokines to being markers of an inflammatory state and there is little information on different functions of the individual cytokines. Therefore, it is not clear to what extent the results of Conejero et al. (2019) regarding IL-1 beta and IL-2 reflect comparable mechanisms as the findings in the present study and why the two previous studies found no effects for IL-6, contradicting our study. Differences in experimental design (social exclusion paradigms vs. social stress) and sample composition (young adults in the study by Slavich et al. and a heterogeneous sample comprising women with a history of suicide attempt, with a major depressive episode without suicidal acts as well as healthy subjects in the study by Conejero et al.) may have played a role here. Also, low statistical power of all studies probably accounts for some of the incoherencies. In the present study, only small clusters of voxels survived rigorous correction for statistical significance and therefore also should be treated with caution. Future studies with high statistical power are needed, which also allow to detect small effects and possibly explain the inconsistent results.

Notably, our finding of a negative association between IL-6 levels and neural activity is in contrast to the theory of Eisenberger and colleagues who assumed that inflammatory markers would correlate positively to neural reactivity underlying social evaluative stress (Eisenberger et al., 2017). However, the theory does not distinguish between acute inflammatory responses and basal circulating levels of cytokines and is mostly based on studies focusing on the effect of brief increases of IL-6 or other cytokines after acute stress induction or as a response to an immune stimulant. In these studies, the association between peripheral inflammation and neural activity of the amygdala, anterior insula, and ACC underlying social processes such as exclusion, rejection, or social pain was positive (Eisenberger et al., 2009; Inagaki et al., 2015; Muscatell et al., 2016, 2015; Slavich et al., 2010). The present findings of an inverted association between basal IL-6 levels and neural activity may thus appear contradictory at first glance. However, it should be noted that the release of IL-6 during acute stress occurs primarily through short-term, stress-induced activation of the sympathetic nervous system (SNS) (Slavich and Irwin, 2014). Likewise, neural (ACC, insula) and physiological (pupil measures) markers of acute stress (e.g., processing of negative emotions or pain) have also been associated with the sympathetic nervous system (Muscatell et al., 2015). Thus, a positive association between stress task-induced increase in IL-6 levels and neural activity can be expected, reflecting individual differences in the stress response to the task. In contrast, in the current study, circulating baseline levels of IL-6 were investigated. Although the sympathetic nervous system is also involved in basal inflammatory processes (Pongratz and Straub, 2014), we assume that the hypothalamus-pituitary-adrenal cortex axis (HPA axis) is another important player in this process. In conditions of elevated levels of inflammation, for example due to lifestyle factors (overweight, chronic stress, alcohol etc.), the complex signaling loops between the brain, the HPA axis and inflammatory processes could be altered (Haroon et al., 2012). To conclude, Eisenberger’s theory on heightened neural reactivity to threats may apply to acute immune processes, whereas basal inflammatory physiology may exert a different effect on neural social stress responses. In this context, however, the possibility that basal levels of IL-6 do not necessarily reflect inflammation must also be considered. In previous research, moderately elevated levels of IL-6 are usually interpreted to reflect an inflammatory state (e.g., Conejero et al., 2019). However, IL-6 has been shown to also have some antiinflammatory properties and IL-6 levels may be elevated in the complete absence of an inflammatory state (Del Giudice and Gangestad, 2018; Scheller et al., 2011). Thus, a broader view of the role of IL-6 has recently been called for (Del Giudice and Gangestad, 2018) and future research should take this into account when investigating interactions with neural processes.

The present findings should be interpreted in the light of some important limitations. First, more than half of the participants did not show an increase in cortisol after the stress task. The percentage of cortisol responders is in line with previous studies using the MIST (Noack et al., 2019), but it is not clear why it is that low. One may speculate that non-responders were not stressed by the MIST, which was, however, not the case in the present study (see S6). There are other possible explanations, which could underlie the lack of cortisol response to the task, for example an anticipatory cortisol response because the scanner environment itself was perceived as stressful (Gossett et al., 2018). The measurement of other stress parameters such as heart rate or skin conductance would be desirable to complement the cortisol results in future studies because more reliable changes by the MIST could be detected in these (Noack et al., 2019). Furthermore, these measures are not as strongly influenced by the circadian rhythm as cortisol. In the present study, it was unfortunately not possible to perform the fMRI measurements at the same time of day, which is why a quantitative evaluation of the cortisol change was not performed. Quantitatively comparable cortisol data and information on awakening cortisol would be important for future studies to be able to better account for HPA axis activity as a possible influencing factor. Second, as the MIST involves higher-order cognitive functions, stress may not be clearly distinguishable from cognitive effort. Also, previous research has shown an inverse relationship between circulating levels of IL-6 and cognitive performance (Marsland et al., 2006). Thus, the present results might not only reflect an association of cytokines with threat-related neural activity, but also with recruitment of neural resources during cognitive task performance. Third, in view of highly discrepant findings on plasma cytokine levels, we focused only on IL-6 which is reliably and frequently measured (Hänsel et al., 2010; Marsland et al., 2017) and usually interpreted as a surrogate for underlying inflammatory processes. However, it can not be considered an unambiguous inflammatory marker (Del Giudice and Gangestad, 2018) and cytokines function as closely coordinated network with pleiotropic, activating, synergistic, antagonizing, and redundant effects (Himmerich et al., 2019). Thus, it would be important to consider more cytokines and additional markers of inflammation in future research to better understand mechanisms of cytokine-brain interactions.

In conclusion, the present study found a negative association between circulating basal IL-6 levels in healthy participants and activity in the AI and amygdala in response to a social evaluative stress task. The results extend on the existing literature on the association between acute immune and neural responses to social threats but indicate an opposite correlation for baseline levels of IL-6. This finding suggests that previous theories about the relationship between immune parameters and brain responses may be too simplistic. For future models of cytokine-brain interactions, it would be desirable to consider both, processes of acute activation of the immune system as well as basal levels of cytokines. In this context, a broader perspective that is not simply treating cytokines as markers of an inflammatory state is demanded. This is of particular relevance, as a disruption of feedback loops between the immune system and the brain is suggested to contribute to the development of psychiatric illnesses (Kraynak et al., 2018).

## Acknowledgments

We would like to thank Christine Klein for her contribution in planning the project, Birte Hell and Nadine Merg for their help with the laboratory work, and Nora Czekalla for help with the analysis of fMRI data.

