## Supplemental material for "Association of stress-related neural activity and baseline interleukin-6 plasma levels in healthy adults"

**Supplementary material**

**S1 Demographic information**

| Characteristic | Total sample (N=65) |
| --- | --- |
| Sex | 60% (N=39) female |
| Age (mean, SD) | 42.6±7.47 years |
| BMI (mean, SD) | 26.4 ±4.95 kg/m^2^ |
| Education | 96.9% (N=63) completed vocational training: N=27 apprenticeship, N=12 technical college, N=8 college, N=15 university, N=1 other |
| Employment | 58.5% (N= 38) full time employed, 27.69% (N=18) part time employed, 13.8% (N=9) unemployed |

**S2 Missing data**

Two participants did not completely fill out the Positive and Negative Affect Schedule (PANAS) at either of the two time points and were thus excluded from the analyses of affective stress experience (difference between the two time points), resulting in a sample of N= 63 for these analyses. One participant did not complete the questionnaires Trier Inventory for Chronic Stress (TICS) and Stress Reactivity Scale (SRS) resulting in a sample of N=64 for analyses involving these questionnaires.

**S3 Montreal Imaging Stress Task (MIST)**

Example of the screen shown during the MIST: 1) an arithmetic task to be solved by participants (left) with possible response options (right); 2) after selecting the answer a feedback is given (on the bottom: “Falsch!” = Wrong!). 3) The upper bar shows the participant’s performance (arrow in red) and the performance of a reference group (arrow in blue); 4) the bar below indicates the elapsed response time.

**S4 fMRI results: stress task-related activation**

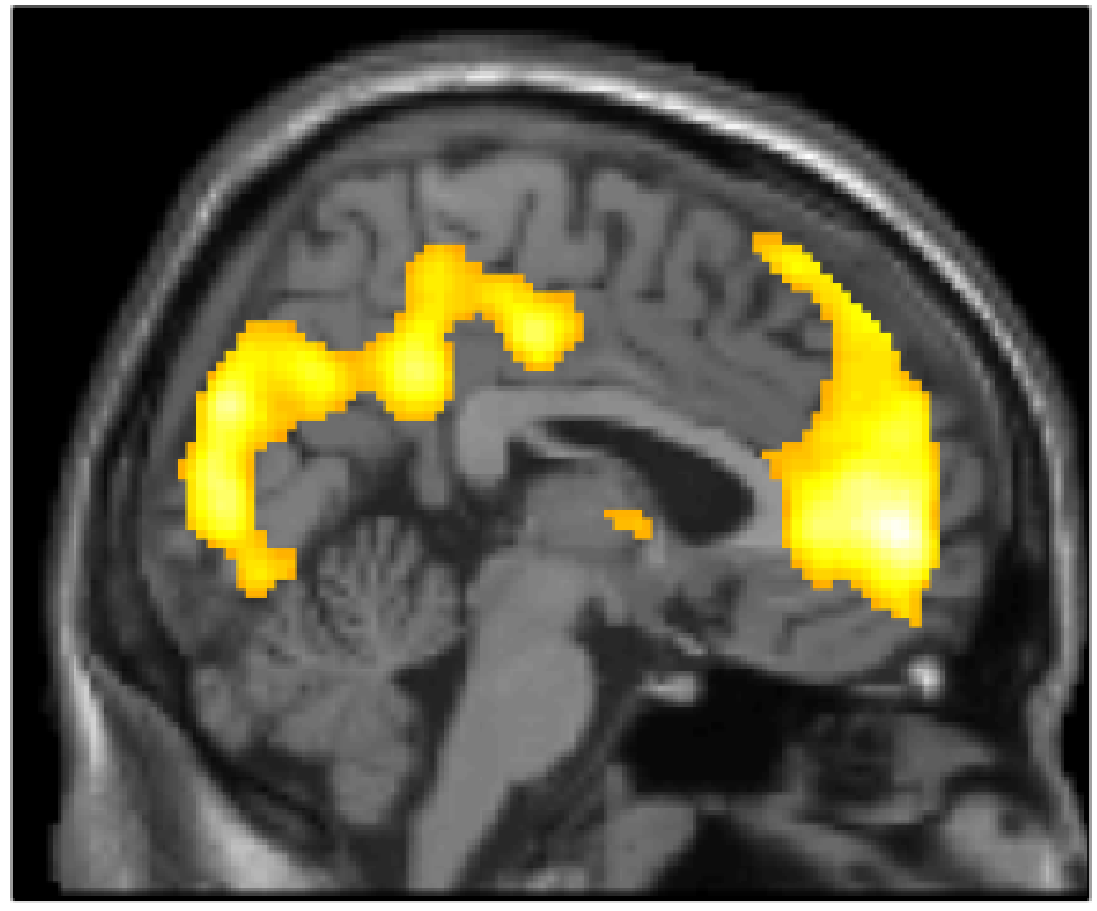

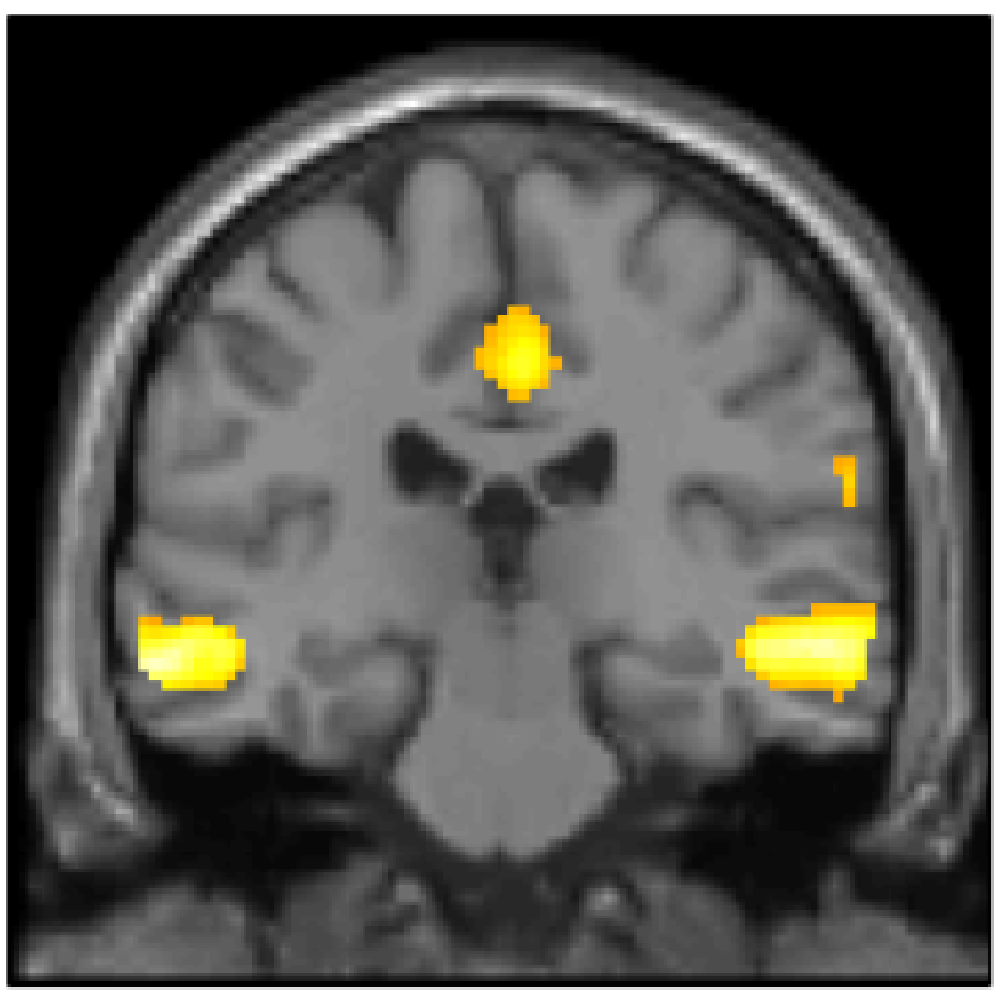

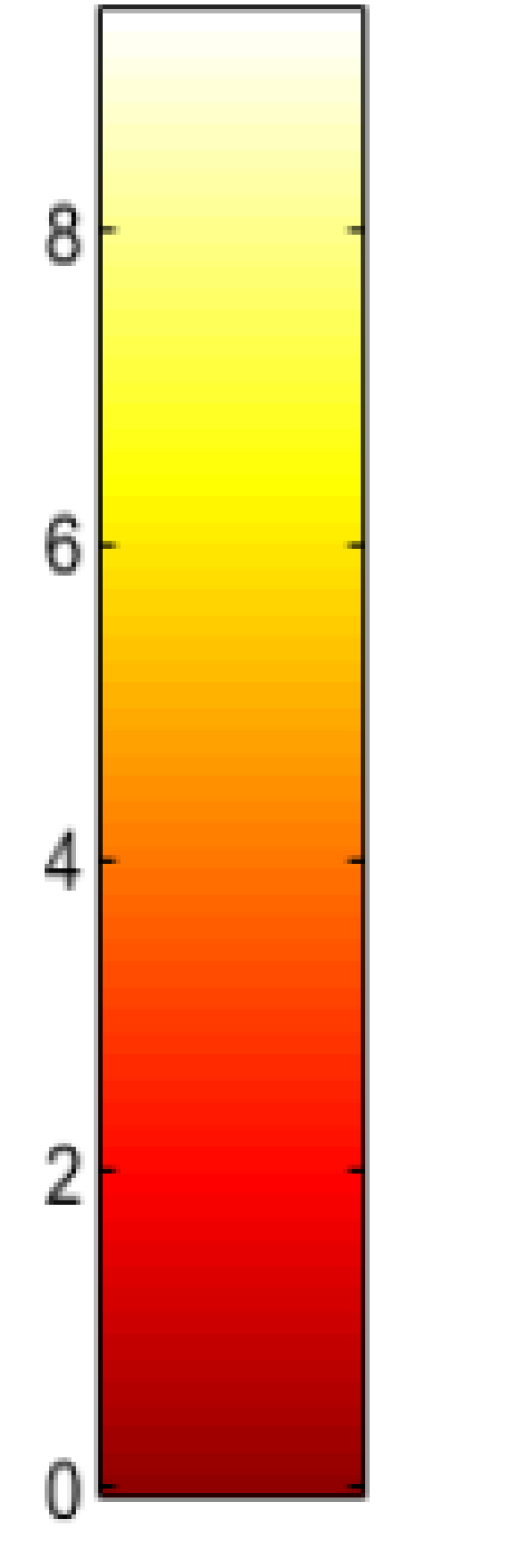

Visualization of the brain activity for the contrast: stress > control condition (p=0.05, FWE corrected).

**stress task-related deactivation**

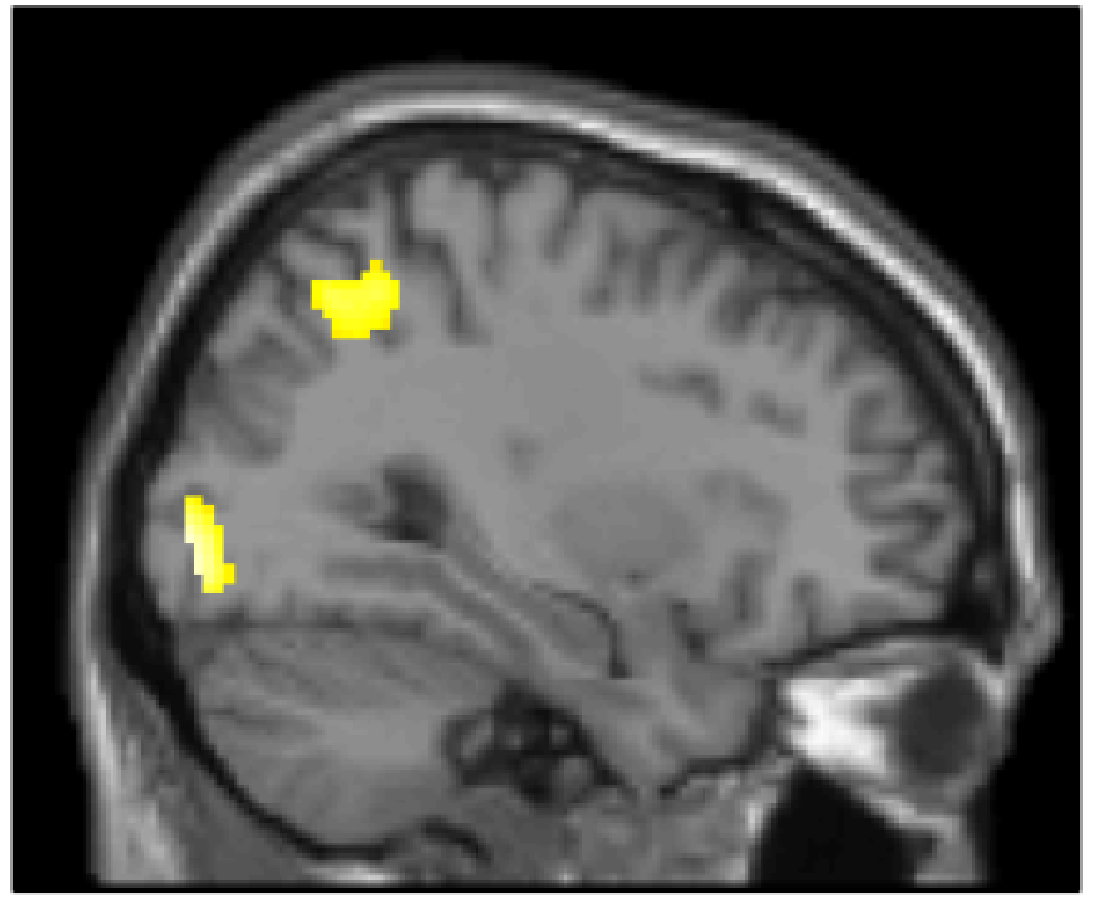

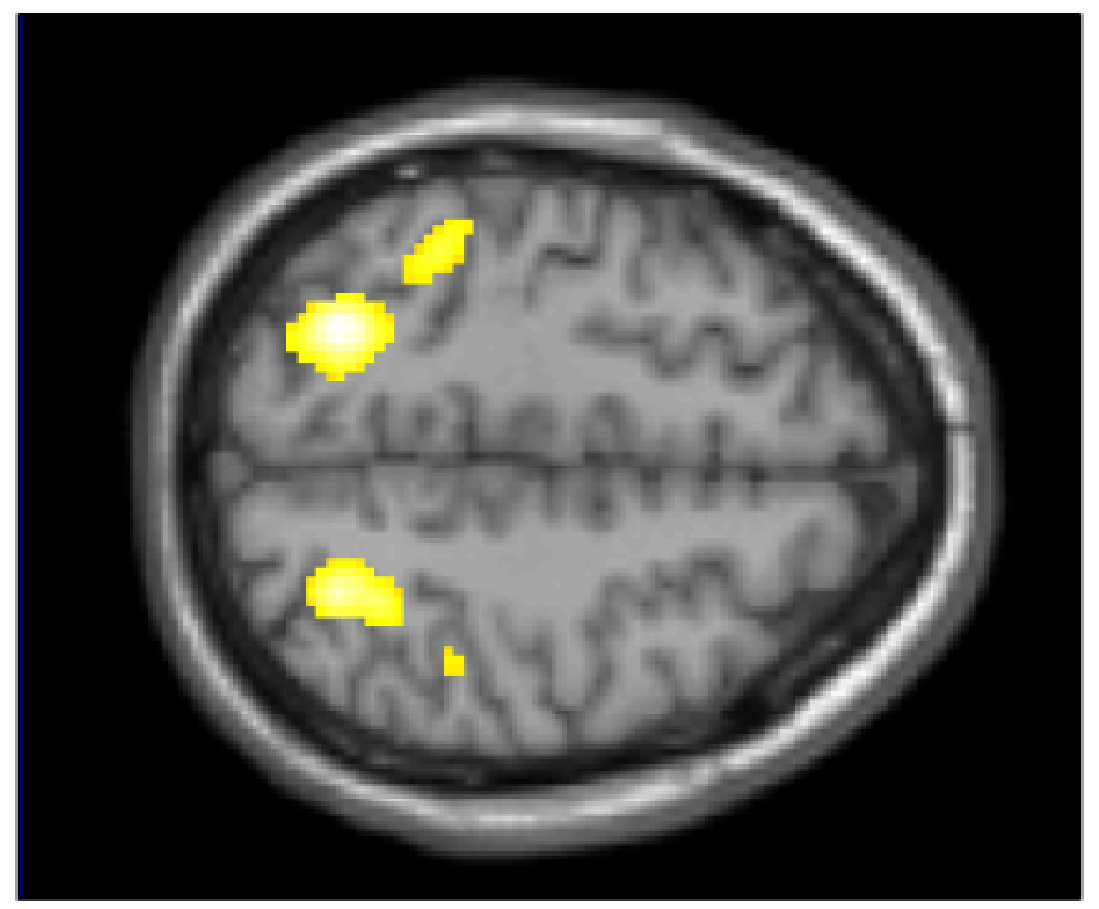

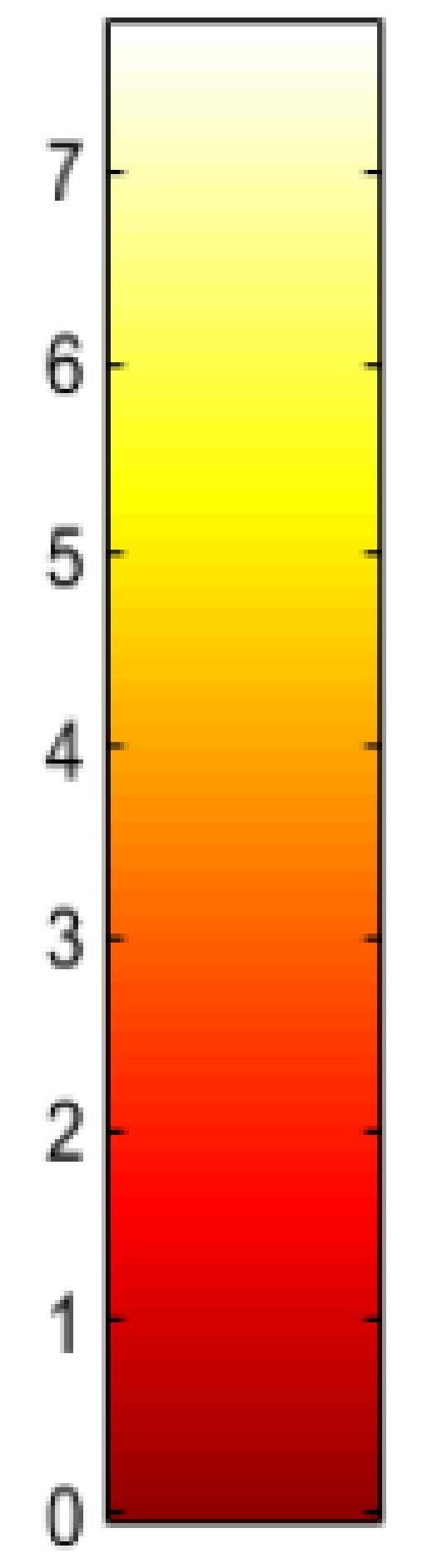

Visualization of the brain activity for the contrast: control condition > stress (p=0.05, FWE corrected).

**S5. Correlations of IL-6 with neural activation at contrast stress > control condition in whole brain analysis (p<0.001, uncorrected).**

|  | side | MNI coordinates | | | Cluster size | T | p (peak) |
| --- | --- | --- | --- | --- | --- | --- | --- |
|  |  | x | y | z |  |  |  |
| *Negative correlation* | | | | | | | |
| Sup. temp. pole | L | -42 | 4 | -20 | 126 | 4.29 | 0.253 |
| Insula | L | -38 | 0 | 10 | 66 | 3.82 | 0.647 |
| Post. cingulum | L | -16 | -36 | 24 | 19 | 3.59 | 0.84 |
| Middle temp. gyrus | L | -48 | 0 | -28 | 15 | 3.58 | 0.849 |
| Middle temp. gyrus | R | 48 | -2 | -16 | 9 | 3.53 | 0.879 |
| Hippocampus | R | 36 | -6 | -26 | 5 | 3.52 | 0.889 |
| Insula | R | 30 | 18 | -18 | 7 | 3.44 | 0.928 |
| Insula | L | -32 | -26 | 26 | 5 | 3.43 | 0.932 |
| Hippocampus | L | -34 | -14 | -18 | 9 | 3.37 | 0.954 |
| *Positive correlation* | | | | | | | |
| Sup. parietal gyrus | R | 20 | -58 | 52 | 6 | 3.4 | 0.943 |

**S6. PANAS results of the group of cortisol non-responder**

To test whether the absence of a cortisol increase in response to the MIST was accompanied by the absence of an affective response, paired t-tests for negative and positive affect (before to after stress induction) were calculated for the subgroup of cortisol non-responders. These showed significant changes in positive (*t*(35) = 4.69, *p* < 0.001) as well as negative affect (*t*(35) = -2.46, *p* = 0.02) from before to after stress exposure, suggesting that they were subjectively stressed by the task, although they did not show a cortisol response.
